# Gaming Addiction Transmission: Interpersonal Neural Pathways Revealed by HYPER-NESS

**DOI:** 10.64898/2026.06.18.732549

**Authors:** Chuyang Sun, Mattia Rosso, Ruoyu Niu, Xinyu Ye, Tinghao Tang, Peter Vuust, Leonardo Bonetti, Rixin Tang

## Abstract

Gaming addiction may be contagious through social interaction. In team competitive video games, individuals of varying addiction levels are often paired. This raises the question of whether, and through what mechanisms, one individual’s addiction level is affected by other players. Methodologically, addressing this question requires understanding both what happens between brains and within individual brains during a gaming session. To this end, we used HYPER-NESS (Hyper Brain Network Estimation via Source Separation), a novel analytical framework to decompose and weight the distinct contributions of inter-brain and intra-brain processes to social brain networks. We applied this framework to a hyperscanning fNIRS dataset where dyads were scanned while playing a video game together against experimenters. We found that inter-brain and intra-brain contributions to the social brain networks were differentially associated with changes in game reward and social reward sensitivity, depending on the addiction level of one’s partner. Granger causality analysis of both social and individual brain networks revealed influences from the high-addiction partner to low-addiction partner. Furthermore, significant cross-frequency coupling was found selectively between low-addiction players, supporting the idea that this form of inter-brain interactions underpins the joint processing of task-relevant rewards. To our knowledge, these findings provide the first neural-level account for the hypothesis that gaming addiction propagates though dyadic social interaction.

## 1. Introduction

With the rapid expansion of the video gaming industry, gaming addiction has emerged as a major global public health concern, with worldwide prevalence estimates ranging from 3% to 5% (Stevens et al., 2021; Limone et al., 2023). Among diverse gaming genres, team competitive games—particularly Multiplayer Online Battle Arena (MOBA) games—exhibit particularly elevated addiction risk profiles (Nuyens et al., 2016; T’ng et al., 2023). One distinctive feature of team competitive games lies in their dual reward environment: players can obtain both game rewards (e.g., kills, victories, skill advancement) and social rewards (e.g., cooperation with teammates, social bonding, relatedness satisfaction) (Przybylski et al., 2010). More importantly, another distinctive feature lies in their heterogeneous team matching environment: random matchmaking mechanisms creating naturalistic conditions under which low-addiction individuals are recurrently exposed to partners whose addiction profiles may differ substantially from their own. Such social dynamic raises the question of whether, and through what mechanisms, such exposure shapes individual reward processing.

Prior theoretical models suggest two pathways through which addiction might spread in such interactive environments. The first pathway proposes a “contagion” process — transmission from high-addiction to low-addiction individuals — which aligns with epidemiological transmission models in complex networks (Pastor-Satorras et al., 2015). We propose that this contagion process may operate through inter-brain neural coordination: coupling with a high-addiction partner may provide a channel through which addiction-related reward-processing patterns gradually propagate onto the low-addiction individual, reshaping their reward sensitivity toward the high-addiction profile. The second pathway proposes “emergence” — spontaneous and collective enhancement of addiction-related motivation among low-addiction individuals through mutual interaction (Jackson et al., 2017; Smaldino, 2023). In this pathway, inter-brain coordination and individual reward processing are expected to mutually reinforce each other symmetrically, such that coordinating with a like-minded partner amplifies shared sensitivity to game-relevant reward signals rather than altering the profile of reward sensitivity itself.

Critically, although these two pathways may not necessarily differ in the degree of addiction risk they confer, we propose that they differ fundamentally in two respects. First, they differ in the type of reward system to which inter-brain coordination is functionally linked: the *social contagion pathway* predicts that stronger inter-brain coupling will be associated with reduced social reward sensitivity — a hallmark of the high-addiction profile toward which the low-addiction individual is converging — whereas the *content resonance pathway* predicts that stronger inter-brain coupling will be associated with enhanced game reward sensitivity through mutual reinforcement. Second, at the timescale of reward-event dynamics, the two pathways predict different causal architectures between individual neural processing and inter-brain coordination: in the *content resonance pathway*, internally generated reward-related neural states in low-addiction individuals are expected to actively drive the formation of interpersonal coordination, whereas in the *social contagion pathway*, this proactive individual-to-dyad influence is expected to be absent, with the inter-brain channel instead reflecting passive propagation from the high-addiction partner (**Figure 1a**.).

**Figure 1.**
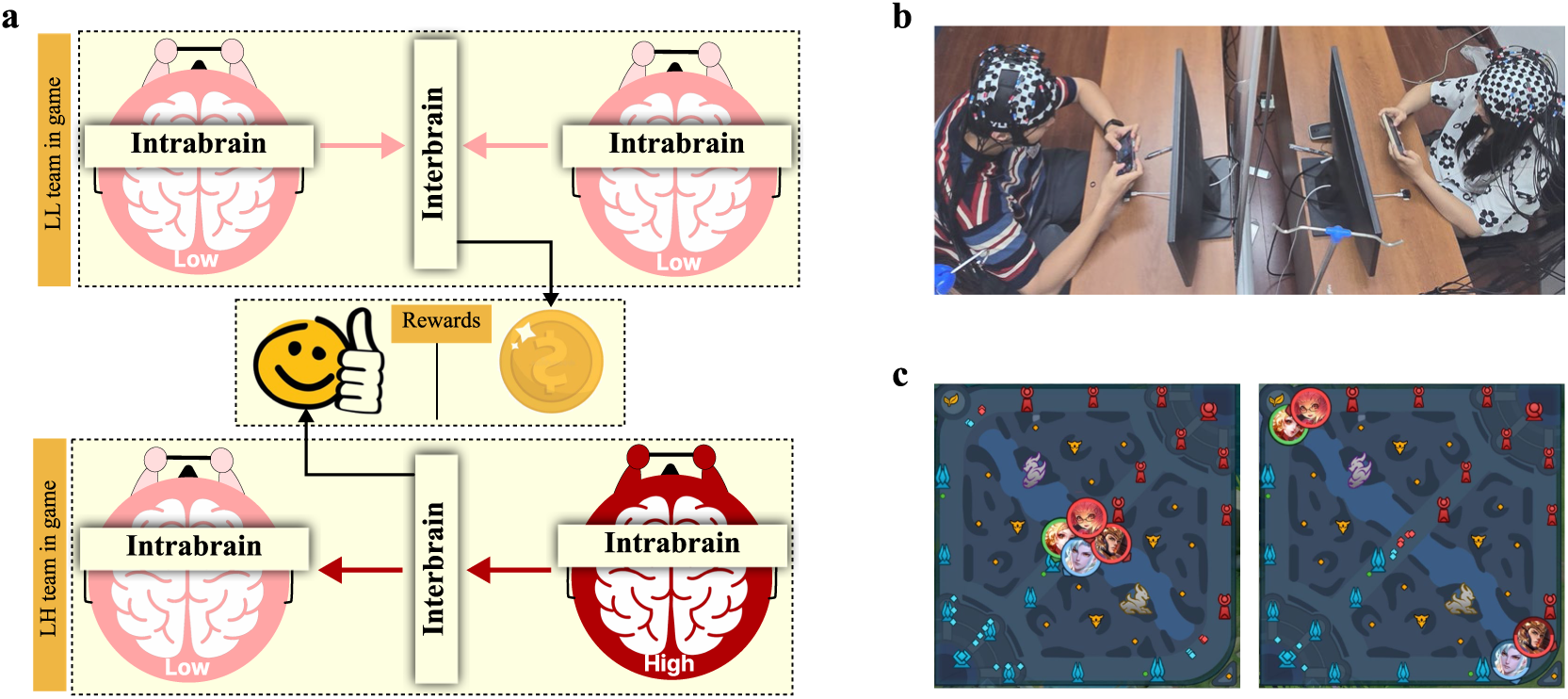
Hypothesized Dual-pathway model of gaming addiction transmission, experimental setup and gaming illustration. **(a)** Upper panel illustrates the *content resonance pathway* (low-low addiction, LL dyads), in which intra-brain contributions from both low-addiction individuals converge symmetrically onto the inter-brain channel, and inter-brain coordination is functionally linked to game reward processing (coin icon). Lower panel illustrates the low-high addiction (LH) dyads, in which neural dynamics originate from the high-addiction individual’s intra-brain contribution (dark red), propagate unidirectionally through the inter-brain channel, and influence the low-addiction partner’s intra-brain processing; here, inter-brain coordination is functionally linked to social reward processing (smiley icon). Arrows indicate the causality. LL = low-addiction dyad; LH = mixed dyad (one low-, one high-addiction individual), **(b)** Photo of the experimental setup, the subjects could not see each other. To protect the privacy of the participants, both individuals in the image are co-authors of the article, and their consent has been obtained. **(c)** Characters’ locations in group fight (left) and single fight (right): the characters not in the red circles are the characters controlled by the participants, and the red circles are the characters controlled by the experimenters. In group fight, the avatars representing the positions of the participants and the experimenters overlap, meaning that the characters controlled by the two participants and the two characters controlled by the experimenters are in the same game scenario. In single fight, the characters controlled by the two participants are competing against one experimenter player in different game scenarios respectively. The avatars representing their positions are located in the upper left and the lower right.

However, quantifying separated inter-brain and intra-brain contributions to reward processing poses a major methodological challenge in social neuroscience. Although the hyperscanning paradigm (i.e., the simultaneous neuroimaging of interacting individuals) has advanced our understanding of the neural correlates of social interaction by revealing inter-brain synchronization during cooperation and coordination (Czeszumski et al., 2020; Kingsbury & Hong, 2020), traditional inter-brain synchronization measures inherently confound genuinely social processes with simultaneous individual responses to shared stimuli. That is, both individuals may show synchronized neural activity simply because they respond to the same game events, not because one brain is influencing the other (Burgess, 2013; Hamilton, 2021). On the other hand, conventional single-brain analyses characterizing neural activity at the level of intra-brain networks’ activation, do not situate individual contributions within the dual-brain system, thereby failing to quantify the relative weight of inter-brain and intra-brain processes. Without separating these two contributions, it is impossible to determine whether reward sensitivity changes arise from inter-brain coordination, intra-brain processing, or their interaction, which makes it impossible to distinguish the two pathways.

Here, we present HYPER-NESS (Hyper Brain Network Estimation via Source Separation) as an extension of the FREQ-NESS framework (Rosso et al., 2025a) to dual-brain hyperscanning scenarios, designed to separate interbrain and intrabrain contributions in the context of social interactions. The mathematical foundation of HYPER-NESS builds upon generalized eigenvalue decomposition (GED) (Cohen, 2022; Rosso et al., 2025a), optimized to identify frequency-specific networks encompassing multiple brains engaged in an interaction. The core principle of FREQ-NESS is to design spatial filters that maximize the contrast between narrowband (target frequency) and broadband (reference) dynamics in a multivariate dataset. This enables the isolation of neural components selectively active at frequencies of interest, while simultaneously characterizing their spatial organization without imposing predefined regions of interest. To this date, this pipeline has been successfully applied in electrophysiology to characterize the dynamic reconfiguration of frequency-resolved brain networks during naturalistic stimulation (Rosso et al., 2025a), following intake of psychedelics (Shinozuka et al., 2025; Shinozuka et al., 2026), and as a consequence of ageing (Malvaso et al., 2025).

The key innovation of HYPER-NESS lies in treating dual-brain data as a unified high-dimensional space. By performing GED on a joint covariance matrix constructed from both participants’ signals simultaneously, HYPER-NESS preserves the inter-brain cross-covariance structure overlooked by intra-brain approaches. At the same time, it moves beyond conventional hyperscanning analysis methods based on interbrain synchrony, which is knowingly a mechanistically opaque construct (Burgess, 2013; Holroyd, 2022; Rosso et al., 2022; Rosso et al., 2025b), modelling interaction as a system of coupled brain networks. This framework enables the decomposition of the total neural variance into intra-brain contributions from each participant and an inter-brain contribution from the cross-covariance across interacting participants— all expressed in the same units and directly comparable within the same analytical framework (for details, see Section 2.4.2.2).

In the context of the present study, our pipeline enables two complementary analyses. First, we can compare the organization of brain networks and their interactions across conditions—examining whether inter-brain and intra-brain contributions differ when playing with high-addiction versus low-addiction partners. Second, specifically for testing the *content resonance path and* the *social contagion path*, we can examine the functional meaning of this balance via correlations with behavioral outcomes. By doing so, we aimed to determine whether inter-brain and intra-brain components show differential relationships with game reward sensitivity as compared to social reward sensitivity. If inter-brain coupling carries different functional significance depending on partner addiction level or task context, this would reveal distinct pathways through which social interaction shapes individual reward processing.

Conceptually, our work draws upon prior cooperative neuroscience research that distinguishes collaborative cooperation (Group Fight, GF) from division-of-labor cooperation (Single Fight, SF) (**Figure 1c**.) (Zhang et al., 2024; Yin et al., 2025). These paradigms differ critically in the functional relationship between social interaction and task completion. Notably, both cooperation types occur naturally within team competitive games, allowing task conditions to be embedded within an ecologically valid gaming context rather than imposed artificially. Especially for collaborative cooperation, interpersonal coordination is a prerequisite for obtaining task rewards, because success depends on moment-to-moment mutual adjustment between partners, which naturally amplifies inter-brain contributions. This distinction provides a framework for designing task conditions that differentially weight inter-brain versus intra-brain contributions.

By integrating the HYPER-NESS methodology for quantitatively separating inter-brain and intra-brain contributions to a social interaction, we address the following core question: What distinct social neural mechanisms do low-addiction individuals exhibit when paired with high-addiction versus low-addiction teammates during team gaming, and how do this neural mechanism differences relate to alterations in reward sensitivity? Specifically, grounded in the dual-pathway model of addiction transmission, we examine: (1) whether inter-brain and intra-brain mechanisms show differential relationships with game reward versus social reward sensitivity; (2) whether these neural-reward relationships differ between low and high addiction; and (3) whether collaborative cooperation and division-of-labor cooperation differ in modulating these patterns. By addressing these questions, we aim to provide neural-level evidence for how social exposure to partners with different addiction levels may shape individual reward processing, thereby elucidating the mechanisms underlying gaming addiction transmission.

## 2. Method

### 2.1 Participants

A priori power analysis with G*Power 3.1.9.6 was performed to determine the sample size in the present study (Faul et al., 2007). Based on our experimental design, at least 34 dyads were required to achieve the expected effect size of 0.25 (Cohen’s f = 0.25) with Type I error probability of 0.05 (α = 0.05) and Type II error probability of 0.05 (1 - β = 0.95) (Cohen, 2013). All subjects were assigned to two different groups: 17 playing with high addicts (LH) groups and 17 playing with low addicts (LL) groups.

The participants’ gaming addiction levels were assessed using the Nine-Item Internet Gaming Disorder Scale (IGD-9) (Lemmens et al., 2015), a comprehensive and widely validated measurement tool (e.g., Ye et al., 2025). Scores of 6 and above were classified as high addiction group, while scores of 4 and below were classified as low addiction group. This resulted in 17 high addiction participants (10 males, 7 females; mean age = 21.41, SD = 2.063) and 51 low addiction participants (24 males, 27 females; mean age = 21.51, SD = 1.984). 68 participants (34 males, 34 females; mean age = 21.49, SD = 1.989) composing 34 dyads were recruited from Nanjing city. To control the impact of gender, the sample included 18 mixed-gender dyads (9 LH groups, 9 LL groups), 8 male-male dyads (5 LH groups, 3 LL groups), and 8 female-female dyads (3 LH groups, 5 LL groups).

To ensure all participants have similar gaming abilities and styles, all participants were recruited from players ranked between “Star Glory” and “King (under 30 stars)” levels in the videogame “Honor of Kings”, with similar play styles and active gaming experience within the past month. Participants did not know each other before the experiment and were prohibited from verbal communication during the study to control for familiarity effects. All participants signed informed consent forms, and both participants in each dyad equally split the total amount of the reward earned in the task.

### 2.2 Task and Procedure

#### 2.2.1 Experimental Task

We employed Honor of Kings (HoK), a widely popular multiplayer online battle arena (MOBA) game with mechanics similar to League of Legends as a game task. In HoK, all players play on a standard map at the same time. The core goal is to improve the team’s overall wins by defeating the opponent’s real-person players in various ways. There are two different ways to win. One way is that multiple real-person players work together to defeat the opponent’s players—which means that the wins are achieved through the cooperation of the roles’ skills (in the current experimental task, we use the common 2 vs. 2 condition occurring in the same place on the map, called Group Fight condition (GF)). The other way is by independent confrontation between one single player and another single (opponent), but still calculate the team’s overall wins. In the current experimental task, we use two simultaneous 1 vs. 1 battles occurring in two different places on the map, called Single Fight condition (SF)) (see **Figure 2**.).

In addition, to create a real gaming context and remove irrelevant variables, we recruited two professional HoK actors as co-experimenters who were trained to flexibly adjust their performance to match the game level of the subjects. Therefore, there were always 4 players in each game, including 2 co-experimenters and 2 subjects. In both the GF condition and the SF condition, each game lasted 360s (minimal time limit for professional HoK actors to surrender) and was played two rounds, so a total of 4 games were played. Each game was manipulated to complete a total of 6 wins of participants. This manipulation resulted in an approximately regular temporal structure of reward events, with six kill events occurring within each 360-second game, corresponding to an average inter-event interval of ∼60 seconds (∼0.017 Hz). Thus, the number of wins and game time were the same in each game.

#### 2.2.2 Experimental Procedure

Two participants who did not know each other came to our laboratory one after another. No communication was allowed in the laboratory. The participants were separated by a partition so that they could not see each other. Each participant sat in front of a laptop which was used to complete the dot probe test. Each of them was assigned an iPhone 16 Pro with HoK installed and logged into an anonymous game account prepared for the experiment. The participants wore the fNIRS device with the help of the experimenter, covering the frontal cortex (FC) and temporoparietal junction (TPJ) area, and they could take it off anytime if they felt any discomfort. After the preparation phase, we measured the baseline reaction time (RT) of each participant with the dot probe paradigm, which indicates the attention bias to social rewards and game rewards (Frewen et al., 2008). Subsequently, they played two consecutive rounds of Group Fight or Single Fight. The total duration of the game task was about 840s (2 rounds of 60s-preparation and 360s game task). Immediately after two rounds of games (GF/SF) ended, we measured once again the participants’ attentional bias using the same dot probe program. The order of Group Fight and Single Fight conditions was pseudo-randomized across dyads.

### 2.3 Data Acquisition

#### 2.3.1 Behavioral Data

Behavioral data were collected using a custom-programmed dot probe task implemented in Python with Tkinter GUI framework. The dot probe paradigm, originally developed by MacLeod et al. (1986) to assess attentional bias toward specific stimuli, was adapted to measure participants’ attention allocation patterns toward game-related visual cues (Lorenz et al., 2013). The experimental procedure consisted of a sequence where participants first viewed a central fixation cross for 500 ms, followed by simultaneous presentation of two images (one game-related reward stimulus and one game-related neutral stimulus; one social-related reward stimulus and one social-related neutral stimulus) for 2000 ms positioned at the left and right sides of the screen. Note that the rewards and neutral stimuli used in the program were selected by experts and are familiar to players. After the stimulus disappeared, a probe (“+”) appeared at either location, and participants were instructed to respond as quickly and accurately as possible by pressing the corresponding arrow key. Each trial was separated by a 1000 ms inter-trial interval. For each trial, the system automatically recorded trial details, probe position, stimulus-probe congruence, response accuracy, and reaction time measured from probe onset to key press.

#### 2.3.2 fNIRS Data

We used a functional near-infrared spectroscopy (LABNIRS, including 780nm, 805nm, and 830nm) from Shimadzu, Japan. Changes in the oxygenated hemoglobin concentration ([HbO]), deoxygenated hemoglobin concentration ([HbR]), and total hemoglobin concentration ([HbT]) in the detection region were obtained according to the modified Beer-Lambert Law (Cope & Delpy, 1988). The sampling rate in this study was 30.3Hz. The self-developed probe distribution was used according to the 10/05 international system (Cutini et al., 2008; Jurcak et al., 2007). There were 20 probes in the measurement area, forming 22 channels (see **supplementary material 1**). The regions of interest (ROI) included bilateral dorsolateral prefrontal cortex (DLPFC), ventrolateral prefrontal cortex (VLPFC), and bilateral temporoparietal junction (TPJ). These regions were covered by four optode probe sets: two 2×3 and two 2×2 configurations, resulting in 22 measurement channels formed by 10 emitters and 10 detectors with 30 mm separation. The spatial anatomical positions of 22 channels were measured by Polhemus Fastrak 3D Digitizer with the virtual spatial registration method, and the corresponding Montreal Neurological Institute (MNI) coordinates were further calculated by the NIRS_SPM MATLAB packages with Nz, Cz, AL, and AR as the referential positions (Singh et al., 2005; Tsuzuki et al., 2007). The location information of all channels covering MNI coordinates and the corresponding Brodmann Area (BA) was provided in supplementary materials.

### 2.4 Data Analysis

#### 2.4.1 Behavioral Data Analysis

Raw data from the dot-probe task underwent systematic preprocessing and analysis. First, the initial trial (practice trial) was removed from each session, followed by retention of only correct response trials, excluding error trials. Subsequently, for each participant, data points exceeding the mean by ± 3 standard deviations were labelled as outliers and removed. Trials were categorized into congruent conditions (probe appeared at the reward location) and incongruent conditions (probe appeared at the neutral stimulus location) based on the relationship between probe position and target stimulus location. Attention bias scores were calculated using the classic dot-probe task analysis method established by MacLeod et al. (1986). The following formula was applied: attention bias score = mean reaction time for incongruent trials − mean reaction time for congruent trials, where positive values indicate attention bias toward reward stimuli and negative values indicate attention avoidance of reward stimuli. Attention bias scores for gaming stimuli and social stimuli were calculated separately for each participant across three experimental conditions (resting baseline, GF condition, SF condition). To quantify the specific effects of different gaming conditions on attention bias, task-induced attention bias change values were further calculated as the difference between GF and SF conditions relative to the resting baseline (GF condition attention bias score − resting baseline attention bias score; SF condition attention bias score − resting baseline attention bias score).

#### 2.4.2 fNIRS Data Analysis

##### 2.4.2.1 Preprocessing on the Participant Level

The preprocessing of fNIRS data was performed in MATLAB (MathWork, Natick, MA, USA) using the Homer2 toolbox. We focused on the HbO data because it is the most sensitive parameter of regional cerebral blood flow and robustly correlates with the BOLD signal of fMRI (Hoshi, 2016). Additionally, HbO has been widely used in previous fNIRS hyperscanning studies on social interactions (Cui et al., 2012; Cheng et al., 2015; Pan et al., 2017; Nozawa et al., 2016).

Data preprocessing followed a systematic series of steps executed in sequence. Initially, signal quality evaluation was performed, with low-quality channels eliminated according to signal-to-noise ratio standards. Channels exhibiting signal-to-noise ratios beneath the threshold of 2 were excluded from further analysis, while detection parameters were configured to 0-3 range and source-detector separation was limited to 0-50 mm to ensure retention of only those channels within acceptable measurement distances. Subsequently, raw light intensity measurements were transformed into optical density values. In the third phase, principal component analysis (PCA) was implemented to eliminate global physiological interference arising from scalp blood flow variations and blood pressure fluctuations (Zhang et al., 2016). PCA was applied to remove global physiological interference, with principal components accounting for up to 80% of the total variance being removed. In the fourth step, the processed optical density data underwent conversion to hemoglobin concentration alterations according to the modified Beer-Lambert law, utilizing partial path-length factors of 6 across all wavelengths. As a final step, head movement artifacts were addressed through the correlation-based signal improvement (CBSI) technique (Cui et al., 2010), which successfully eliminates motion-related artifacts while maintaining the integrity of underlying hemodynamic responses. All datasets satisfied quality criteria with no exclusions from the analytical procedures.

##### 2.4.2.2 HYPER-NESS: Hyper Brain Network Estimation via Source Separation

###### Mathematical Foundation of GED

HYPER-NESS builds upon generalized eigenvalue decomposition (GED), which has been widely adopted in multichannel electrophysiology to design spatial filters to maximize the contrast between broadband and narrowband dynamics of interest (Cohen, 2017, 2022; Rosso et al., 2021, 2022, 2023, 2025; Baliviera et al., 2025; Moumdjian et al., 2025; Zuure & Cohen, 2021). More recently, it was implemented in the FREQ-NESS pipeline to separate frequency-specific brain networks by designing spatial filters that maximize the ratio of narrowband to broadband covariance (Rosso et al., 2025a; Shinozuka et al., 2025; Shinozuka et al., 2026; Malvaso et al., 2025).

For a frequency of interest, given the narrowband covariance matrix **S** and broadband reference covariance matrix **R**, GED solves the following eigenequation:

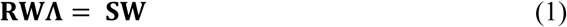

where **W** represents the matrix of eigenvectors (spatial filters) and Λ represents the diagonal matrix of eigenvalues. The eigenvectors identify weighted combinations of channels that best separate frequency-specific oscillatory activity from broadband background activity, while eigenvalues quantify the explained variance ratio for each corresponding signal component.

A critical mathematical property of GED is that the sum of all eigenvalues equals the total variance ratio represented by the decomposition. Specifically, for *k* decomposed components:

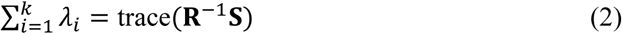

This property ensures the completeness of variance decomposition: the contribution of each component can be precisely quantified, and the sum of all components’ contributions equals the total explainable variance. This mathematical property provides the foundation for HYPER-NESS’s variance decomposition capability.

###### Dual-Brain Covariance Structure

For the combined dual-brain data matrix [44 × T] formed by concatenating both participants’ 22-channel HbO time series, the narrowband covariance matrix exhibits a block structure:

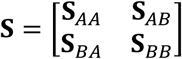

where **S**_AA_ and **S**_BB_ represent intra-brain covariance within each individual, and **S**_AB_ = **S** ^T^ represents inter-brain cross-covariance. Traditional approaches applying GED separately to each participant inherently discard the off-diagonal inter-brain information. HYPER-NESS, on the other hand, preserves this information by performing GED on the complete joint 44×44 matrix, yielding joint eigenvectors **w** that reflect global optimization for frequency-specific dynamics across both brains simultaneously. The joint eigenvector is then partitioned into two 22-dimensional sub-vectors:

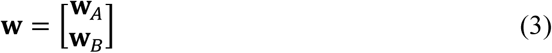

###### Frequency Parameters and Component Selection

For each dyad and task condition, GED was computed over a frequency range of 0.01–1 Hz (resolution: 0.002 Hz), with filter widths of 0.01 Hz for ultra-low frequencies (<0.02 Hz) and 0.02 Hz otherwise. [MR1.1] A regularization shrinkage factor of 0.02 was applied to the R covariance matrix. The top 10 components ranked by eigenvalue magnitude were extracted at each frequency point; subsequent analyses focused on Component 1, which by GED optimization captures the primary frequency-specific network pattern (Cohen, 2022; Rosso et al., 2025a). Group differences in frequency-specific engagement were identified via 2 (Group) × 2 (Task) repeated-measures ANOVAs on Component 1 eigenvalues at each frequency, with cluster-based permutation correction (Maris & Oostenveld, 2007). Frequency bands potentially contaminated by physiological artifacts (cardiac ∼1 Hz, respiration ∼0.2–0.3 Hz) were excluded following established fNIRS artifact ranges (Sasai et al., 2011; Tachtsidis & Scholkmann, 2016).

###### Static Variance Decomposition

For each identified frequency cluster, total variance is decomposed exactly into inter- and intra-brain contributions via reprojection onto the covariance blocks (Haufe et al., 2014):

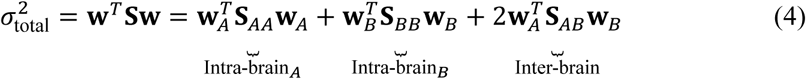

###### Dynamic Time Series Recovery

To examine directed temporal dynamics, network activation time series were recovered by spatially filtering the data through the eigenvectors. For each frequency within significant clusters, raw dual-brain data were narrowband-filtered (zero-phase Butterworth, 4th order, *f* ± 0.002 Hz) and projected onto the GED-derived eigenvectors to yield participant-level component signals S_A_ (t) and S_B_ (t). Instantaneous inter- and intra-brain contribution time series were then computed as:

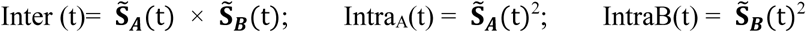

For LH dyads, Intra_L was isolated via predetermined position mapping identifying the low-addiction participant; for LL dyads, Intra_L was averaged across both participants given their symmetric addiction profiles. The temporal mean of each instantaneous series was verified to correspond to its respective quadratic form in Eq. (4) (relative error < 5% across all dyads), confirming analytical consistency between static and dynamic decompositions. These time series were subsequently submitted to Granger causality and cross-frequency coupling analyses.

###### Brain network timeseries interaction

At a later stage of the pipeline, these time series were submitted to two parallel directed analyses. Granger causality (GC) analysis employed bivariate VAR models fit to the [Inter(t), Intra_L(t)] series to quantify directed predictive relationships between inter- and intra-brain contributions; model order selection, permutation scheme, and cross-lag robustness checks are detailed in Appendix A. Cross-frequency coupling (CFC) analysis quantified phase-amplitude coupling between the 0.02 Hz and 0.144 Hz components using the modulation index (Tort et al., 2010); analysis details and cross-subject CFC procedures are provided in Appendix B. For the pipeline, please find it in Figure 3.

**Figure 3.**
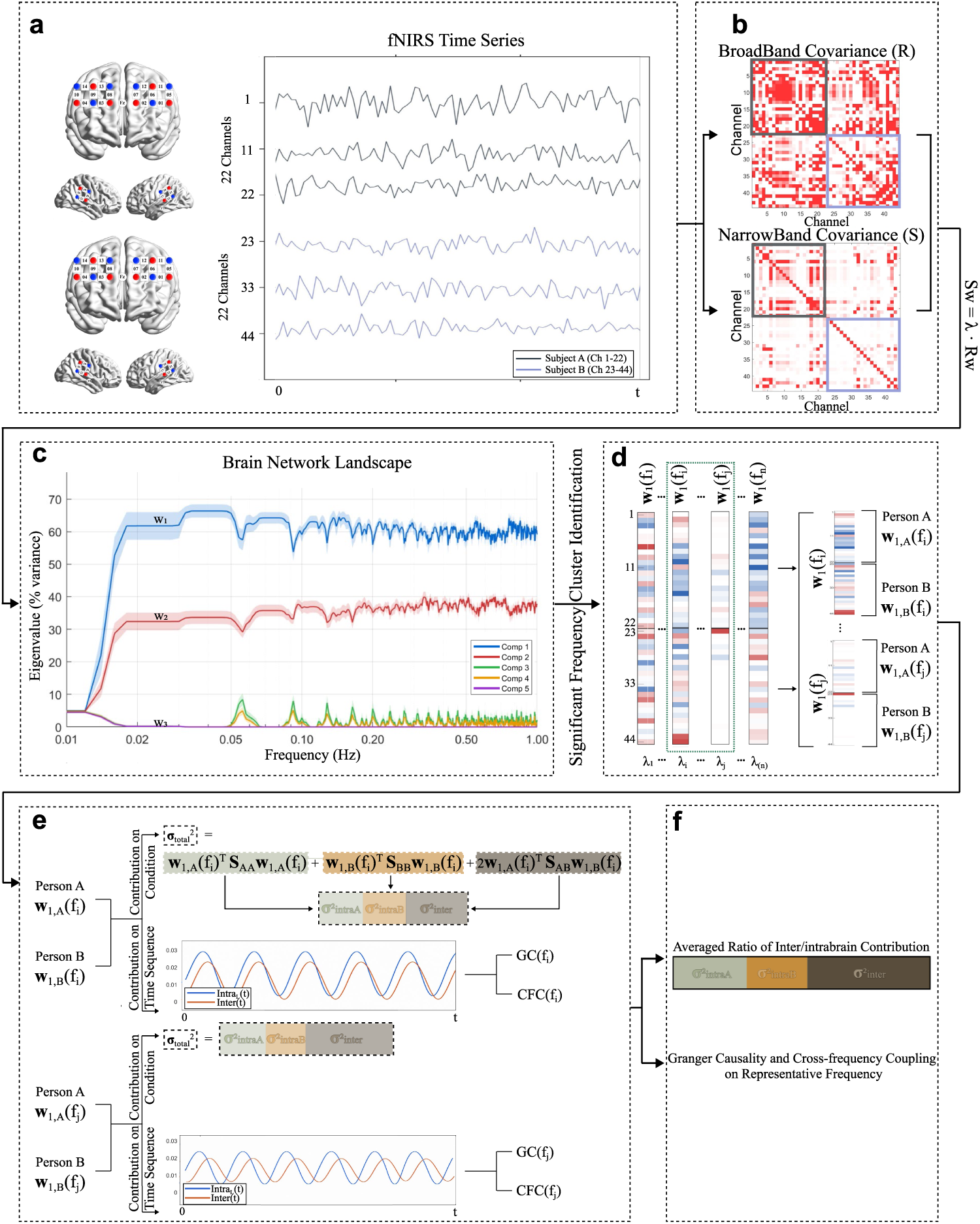
HYPER-NESS analytical pipeline. The panel illustrates our pipeline for decomposing inter-brain and intra-brain neural contributions in dual-brain hyperscanning. **(a)** fNIRS dual-brain hyperscanning time-series were integrated into a unified data matrix (22+22 channels × Time). **(b)** Covariance matrices were computed from the broadband joint data matrix (*R*) and from the narrowband-filtered joint data matrix (*S*); narrow-band filtering was iterated across 0.01 Hz to 1 Hz. Both S and R joint covariance matrices can be partitioned into submatrix blocks: the darker frame indicates the intra-brain covariance matrix of Participant A, the lighter frame indicates the intra-brain covariance matrix of Participant B, and the off-diagonal blocks represent the inter-brain cross-covariance between the two participants. **(c)** Schematic representation of the eigenvectors (*W*) and eigenvalues (*Λ*) obtained through Generalized Eigendecomposition (GED), where the eigenvalues express the variance explained by each network component. **(d)** Partitioning of the joint eigenvector *w* = [*w_A_*; *w_B_*] into individual-specific subvectors for subsequent decomposition of inter-brain and intra-brain contributions in frequency bands. **(e)** At each frequency point within identified clusters, inter- and intra-brain variance contributions were quantified via inner products with the corresponding covariance blocks, and instantaneous contribution time series were recovered from GED spatial filters. These time series were submitted to Granger causality and cross-subject cross-frequency coupling analyses to characterize directed temporal dynamics between inter- and intra-brain contributions. **(f)** Inner product values were averaged across frequency points within each cluster to yield band-level scalar contributions for group comparison and neural-behavioral correlations. Granger causality and cross-frequency coupling results were additionally verified for consistency across all frequency points within each band.

## 3. Results

### 3.1 Eigenvalue Spectrum and Component Selection

Examination of the eigenvalue spectrum revealed a pronounced exponential decay in the variance explained by the GED components across all frequency points. Specifically, Component 1 consistently accounted for the highest proportion of variance (averaged across all frequency points: 59.95% ± 1.87%), with subsequent components showing exponentially diminishing contributions (averaged across frequency points: Component 2: 36.67% ± 1.72%; Component 3: 1.71% ± 0.38%%). Based on this observation, subsequent analyses focused exclusively on Component 1 to reduce data dimensionality while minimizing information loss. Results of Component 2 on frequency points are given in Table S1.

### 3.2 Brain network landscapes

To identify frequency bands exhibiting differential neural patterns between LL and LH groups, we conducted a 2 (Group: LL vs. LH) × 2 (Task: GF vs. SF) repeated-measures ANOVA on Component 1 eigenvalues at each frequency point. Cluster-based (2 frequency points) permutation testing (Maris & Oostenveld, 2007) was applied with 1000 permutations to control for multiple comparisons across frequencies. Frequency bands potentially contaminated by physiological artifacts such as cardiac rhythm (∼1 Hz) and respiration (∼0.2–0.3 Hz) were excluded from the analysis following established fNIRS artifact frequency ranges (Sasai et al., 2011; Tachtsidis & Scholkmann, 2016). Cluster-based permutation testing revealed 2 significant cluster(s) showing Group main effects: 0.02 – 0.026 Hz (4 frequency bins, p = 0.007); 0.144 – 0.148 (3 frequency bins, p = 0.031) (**Figure 5**.).

**Figure 4.**
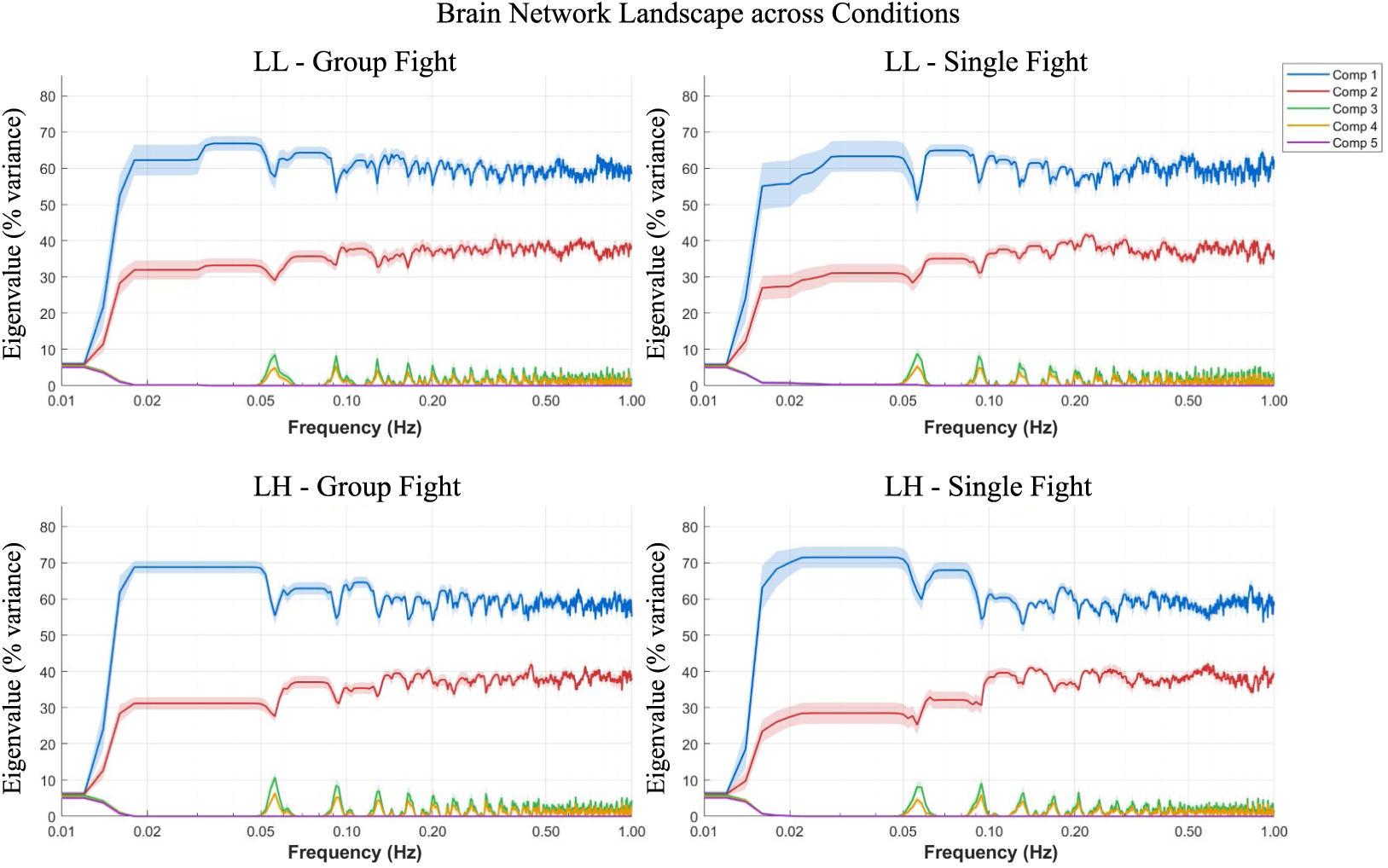
GED eigenvalue spectra across four experimental conditions. Each panel displays the normalized eigenvalue (% variance explained) of the top five GED components as a function of frequency (Hz, log-scaled). Shaded regions indicate ±1 standard error across dyads.

**Figure 5.**
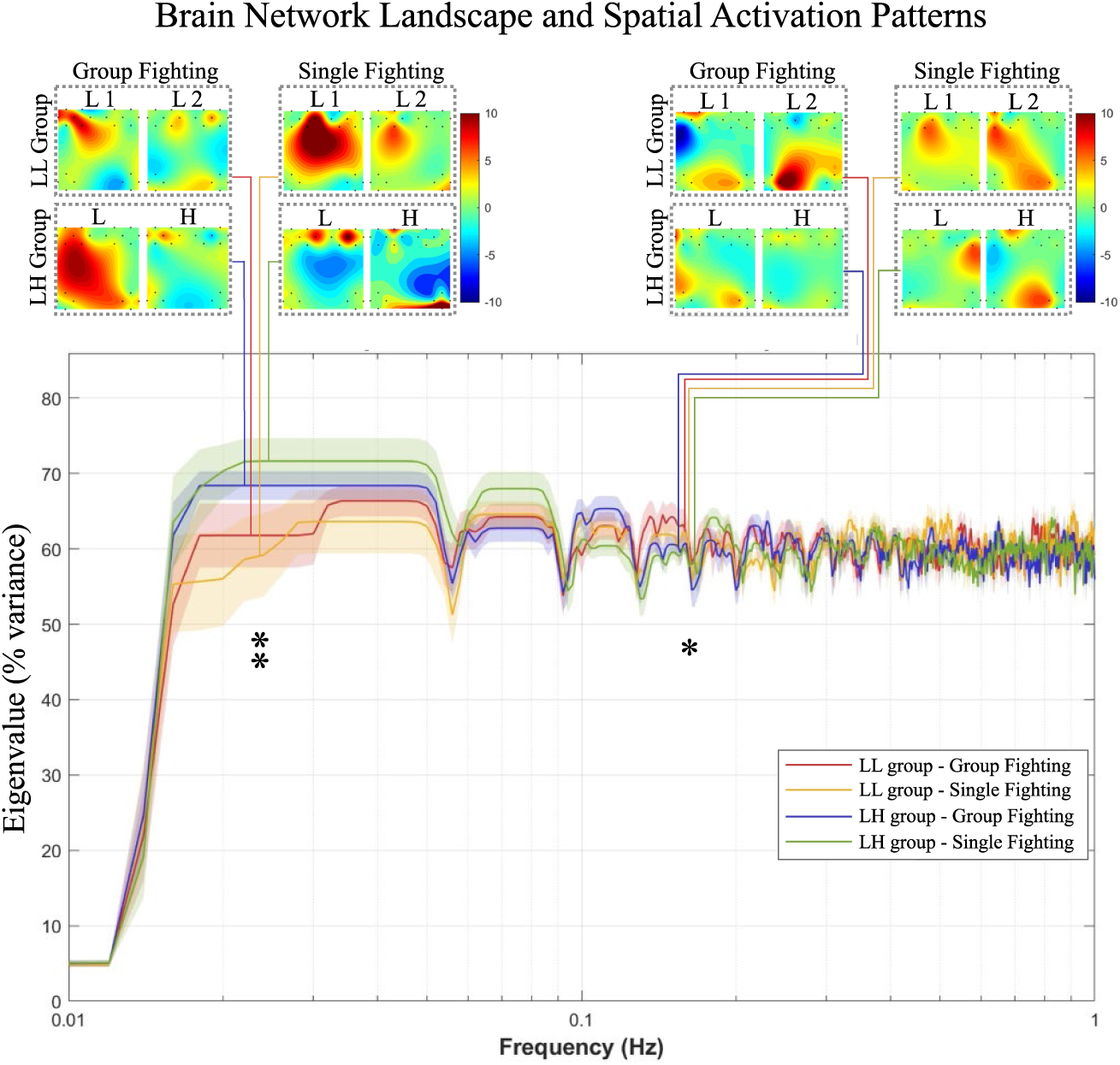
Leading component eigenvalues across frequencies and corresponding spatial activation patterns. The y-axis represents the normalized eigenvalue (% variance explained) of the leading GED component (Component 1), and the x-axis represents frequency in Hz (log-scaled). Lines represent group × task condition means (LL - GF, LL - SF, LH - GF, LH - SF), with shaded regions indicating ±1 standard error. Topographic maps (insets) display the spatial activation patterns of the leading component within each significant frequency cluster, separately for each group (LL and LH) and task condition (GF task and SF task); warmer colors (red) indicate channels contributing more strongly to the leading component, while cooler colors (blue) indicate lower contribution. Vertical lines mark the position of frequency clusters showing significant group main effects as identified by cluster-based permutation testing. Stars denote frequency bins where significant differences were detected.

### 3.3 Variance Decomposition: weighting Inter-Brain and Intra-Brain Contributions

Within the two significant frequency bands, we applied quadratic form analysis to Component 1 to quantitatively separate inter-brain from intra-brain contributions. The magnitude of these contributions was quantified based on the inner product between the eigenvector subcomponents and their corresponding covariance blocks (Eq. 4). Each contribution was then expressed as a proportion of the total variance, yielding inter-brain and intra-brain contribution ratios that served as the dependent variables in subsequent analyses.

#### Frequency Band: 0.02–0.026 Hz

A 2 × 2 (Task: GF task vs. SF task; Group: LL vs. LH) mixed ANOVA on the proportion of inter-brain or intra-brain contributions revealed no significant main effects or interactions (all *p*s > 0.05), indicating comparable contribution magnitudes across experimental conditions in this ultra-low frequency range.

#### Frequency Band: 0.144–0.148 Hz

For intra-brain contribution proportions, we observed a marginally significant Task × Group interaction, *F* (1, 38) = 3.95, *p* = 0.056, η²*p* = 0.094. Post-hoc pairwise comparisons revealed that within the LH group, intra-brain contributions were significantly greater during SF task compared to GF task (*t* = 2.649, *p* = 0.013), suggesting task-dependent modulation of intra-brain network engagement specifically when low-addiction individuals interacted with high-addiction partners (**Figure 6**.).

**Figure 6.**
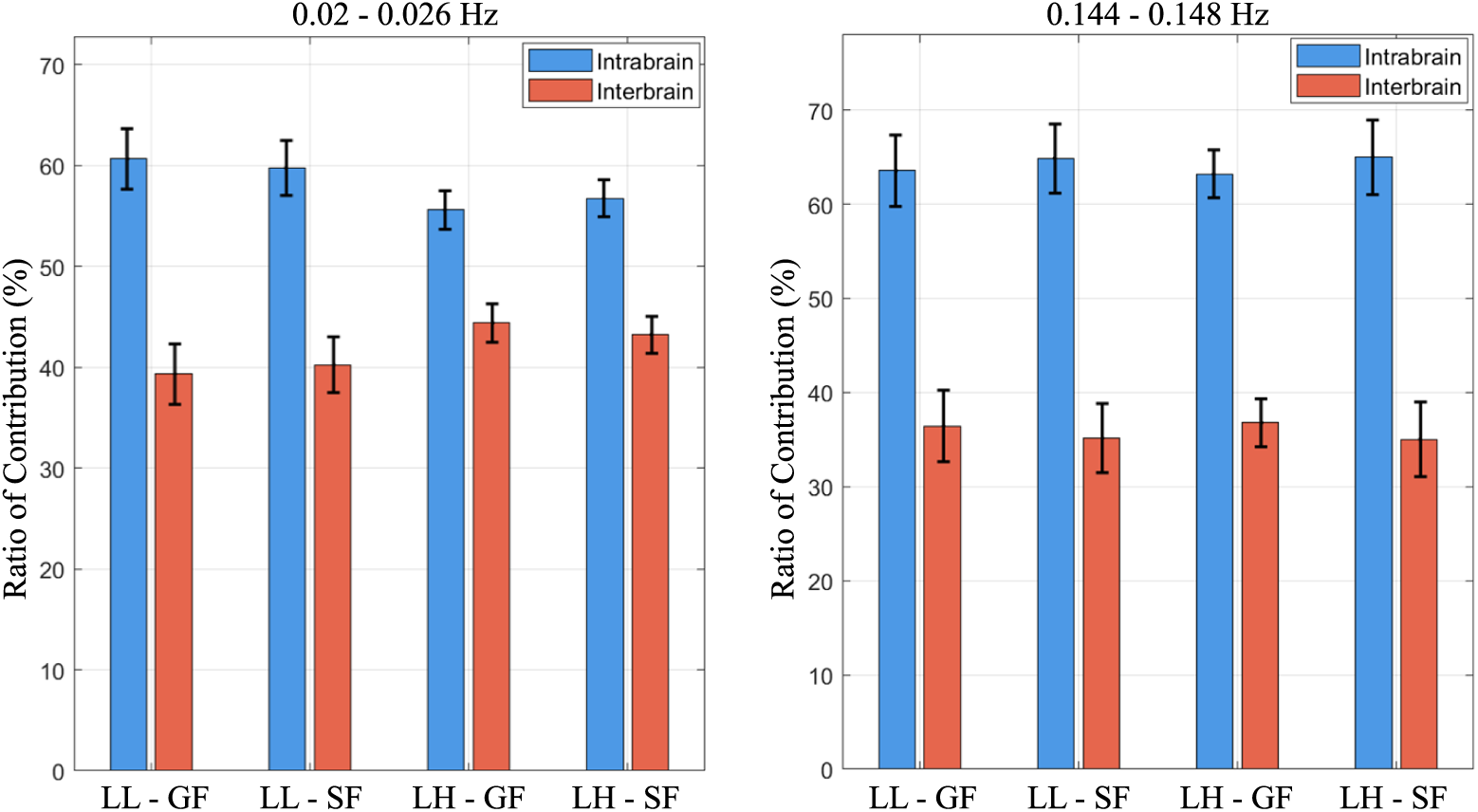
Inter/intra brain contribution ratio to frequency-specific dynamics. Bar plots show the ratio of intra-brain (blue) and inter-brain (orange) contributions to total contributions for LL and LH dyads under Group Fight (GF) and Single Fight (SF) conditions, at 0.02–0.026 Hz (left) and 0.144–0.148 Hz (right). Error bars represent standard error of the mean.

**Figure 7.**
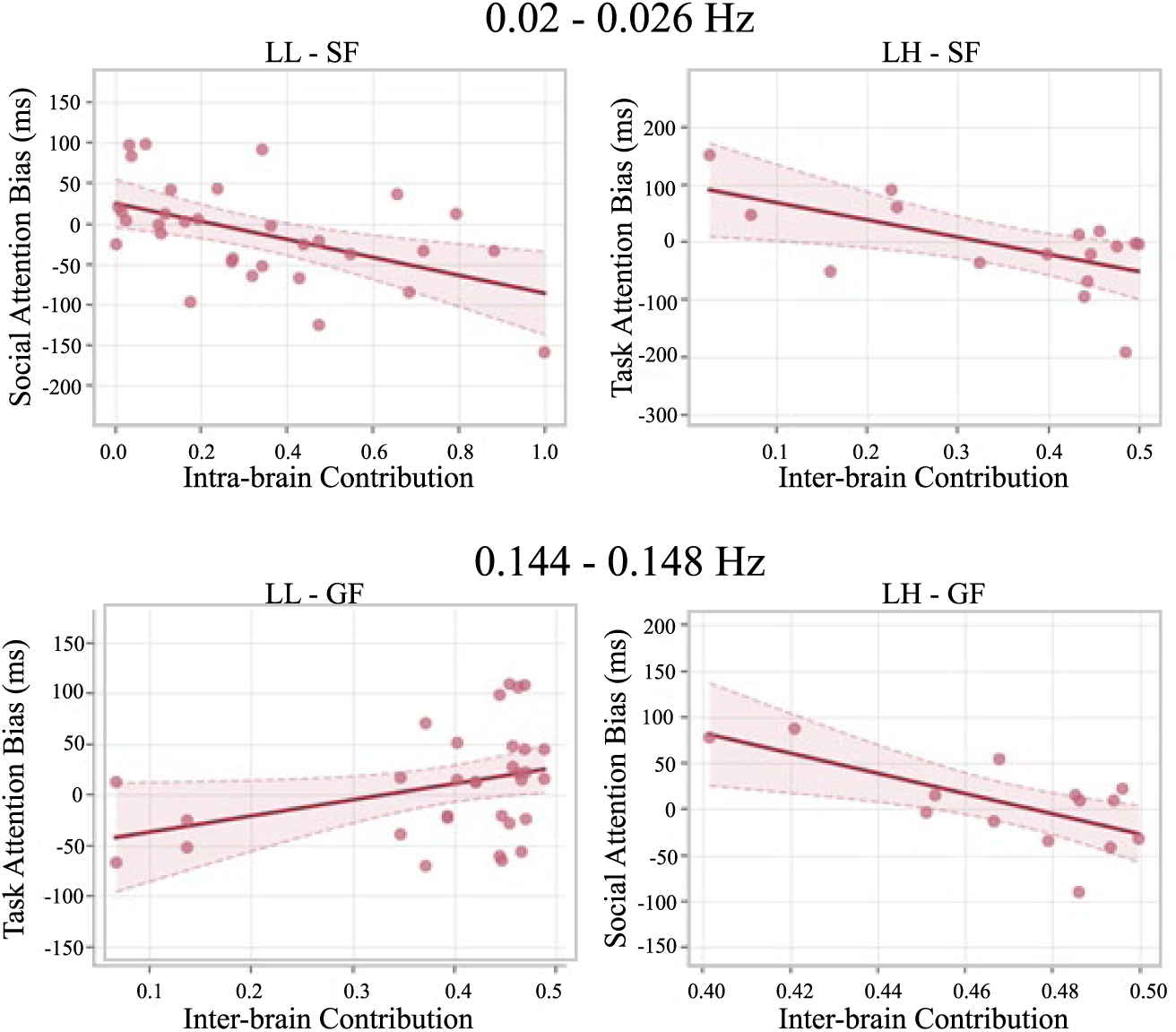
Significant correlations of inter/intra contributions and attention bias. Upper panels show results at 0.02–0.026 Hz: intra-brain contribution negatively correlated with social attention bias in LL dyads during SF task (left), and inter-brain contribution negatively correlated with game reward attention bias in LH dyads during SF task (right). Lower panels show results at 0.144–0.148 Hz: inter-brain contribution positively correlated with game reward attention bias in LL dyads during GF task (left), and inter-brain contribution negatively correlated with social attention bias in LH dyads during GF task (right). Dotted lines indicate 95% confidence intervals.

### 3.4 Neural-Behavioral Correlations

#### Frequency Band: 0.02–0.026 Hz

During SF task, distinct correlation patterns emerged across groups. In LH dyads, low-addiction individuals’ inter-brain contributions were negatively correlated with game reward attention bias (r = -0.603, p = 0.013), suggesting that greater inter-brain coupling was associated with reduced attention capture by game-related stimuli. Conversely, in LL dyads, low-addiction individuals’ intra-brain contributions were negatively correlated with social reward attention bias (r = -0.506, p = 0.003), indicating that stronger intra-brain network engagement corresponded to diminished social attention.

#### Frequency Band: 0.144 – 0.148 Hz

During GF task, we observed group-specific neural-behavioral associations. In LL dyads, low-addiction individuals’ inter-brain contributions were positively correlated with game reward attention bias (r = 0.358, p = 0.044), whereas in LH dyads, their inter-brain contributions were negatively correlated with social reward attention bias (r = -0.676, p = 0.008, n = 14, two outliers removed based on iterative Grubbs test on inter-brain contribution values).

No other significant correlations were observed across the remaining task-group combinations (all ps > 0.05), indicating that identical neural components carry distinct functional meanings depending on the addiction profile of one’s interaction partner.

### 3.5 Granger Causality: Directionality of Inter-Brain and Intra-Brain Coupling

To examine the directionality of inter-brain and intra-brain neural dynamics in driving the interaction, we computed Granger causality (GC) between the Inter and IntraL time series at each frequency point within the two identified clusters (Cluster 1: 0.020–0.026 Hz; Cluster 2: 0.144–0.148 Hz), using a VAR model with lag = 4. For LH dyads, GC was additionally computed between Inter and IntraH to characterize the high-addiction participant’s directional contribution. Cross-lag robustness was confirmed at lag = 3 and lag = 5 (see **Appendix A**). GC values and directional significance patterns were consistent across all frequency points within each band, confirming within-band stability of the observed effects. For Cluster 1, near-identical values across the four frequency points reflect the broad filter bandwidth relative to frequency spacing (FWHM = 0.02 Hz; see Section 2). As the two frequency clusters were identified on the basis of significant group differences in neural dynamics, the following directional patterns are interpreted with particular attention to how GC results differentiate between LL and LH dyads. Results of Granger Causality and their significances are shown in **Table 1a** and **1b**.

**Table 1a.**
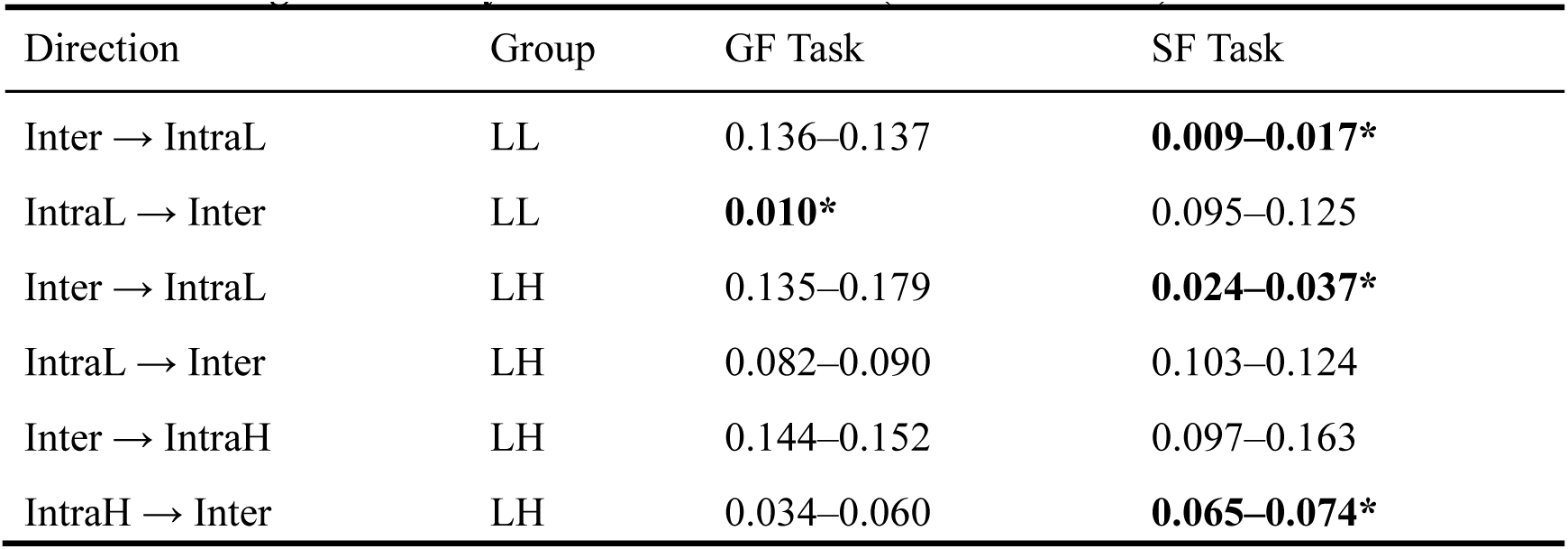
Granger Causality Results on Cluster 1 (0.020–0.026 Hz)

| Direction | Group | GF Task | SF Task |
| --- | --- | --- | --- |
| Inter → IntraL | LL | 0.136–0.137 | **0.009–0.017\*** |
| IntraL → Inter | LL | **0.010\*** | 0.095–0.125 |
| Inter → IntraL | LH | 0.135–0.179 | **0.024–0.037\*** |
| IntraL → Inter | LH | 0.082–0.090 | 0.103–0.124 |
| Inter → IntraH | LH | 0.144–0.152 | 0.097–0.163 |
| IntraH → Inter | LH | 0.034–0.060 | **0.065–0.074\*** |

**Table 1b.** Granger Causality Results on Cluster 2 (0.144–0.148 Hz)

| Direction | Group | GF Task | SF Task |
| --- | --- | --- | --- |
| Inter → IntraL | LL | **0.008–0.018\*** | **0.011–0.020\*** |
| IntraL → Inter | LL | **0.010–0.023\*** | **0.017–0.019\*** |
| Inter → IntraL | LH | **0.031–0.042\*** | **0.011–0.021\*** |
| IntraL → Inter | LH | **0.016–0.023\*** | **0.019–0.023\*** |
| Inter → IntraH | LH | **0.391–0.566\*** | **0.662–0.727\*** |
| IntraH → Inter | LH | **0.191–0.350\*†** | **0.334–0.497\*** |

Note: \* $p < 0.05$ ; \*† marginal significance.

### 3.6 Cross-Frequency Coupling

Building on the functional interpretations established in Section 2 and 4.1—where C1 (∼0.020 Hz) corresponds to the game reward event cycle (∼50 s per kill) and C2 (∼0.144 Hz) reflects a background interpersonal coordination rhythm (Park et al., 2022)—we examined whether these two frequency clusters interact across individuals by computing cross-subject phase-amplitude coupling (PAC) using the modulation index (MI; Tort et al., 2010): C1 phase modulating C2 amplitude (C1ph→C2amp). Robustness across all 12 frequency-pair combinations within the two clusters is reported in **Appendix B**.

For the C1ph→C2amp direction, a significant symmetric cross-subject MI was observed in LL dyads during GF task, indicating that one individual’s slow-wave phase reliably modulated the other’s fast cooperative rhythm. This effect was absent in LL dyads during SF task, and neither the H→L nor the L→H direction reached significance in LH dyads in either task (see **Table 2**). The significant coupling was therefore directionally specific: slow-wave dynamics drove fast cooperative rhythms across individuals.

**Table 2.** Cross-Subject Cross-Frequency Coupling (C1ph→C2amp)

| Group | GF Task | SF Task |
| --- | --- | --- |
| LL | **0.136\*** | 0.129 |
| LH (H→L) | 0.104–0.109 | 0.106–0.107 |
| LH (L→H) | 0.103–0.109 | 0.107–0.108 |

Note: \* $p < 0.05$ .

## 4. Discussion

In this study, we applied HYPER-NESS to decompose simultaneously recorded fNIRS signals during dyadic gaming interactions into inter-brain and intra-brain neural components. This decomposition revealed two frequency bands (∼0.02 Hz and ∼0.144 Hz) that differentiated low-low addiction (LL) and low-high addiction (LH) dyads, suggesting that interaction-related neural dynamics operate across multiple timescales.

These frequency components may correspond to distinct aspects of the task structure. Specifically, the ∼0.02 Hz component aligns with the approximate inter-event interval of game rewards (∼60 seconds), as imposed by the experimental design, whereas the ∼0.144 Hz component falls within a frequency range (0.075 – 0.15 Hz) previously associated with interpersonal coordination in fNIRS hyperscanning research (Park et al., 2022). These interpretations provide a useful framework for organizing the observed neural–behavioral and dynamical patterns, although they should not be taken as a strict mapping between frequency and function. Notably, although the primary focus of the present study is on differences between LL and LH dyads, these differences manifest differently across task contexts (Group Fight vs. Single Fight), suggesting that the observed patterns are modulated by interaction structure rather than being task-invariant.

Within this framework, three complementary analyses — neural – behavioral associations, Granger causality, and cross-frequency coupling—revealed converging but partial patterns that are broadly consistent with the proposed dual-pathway account. Section 4.1 discusses these findings in relation to the dual-pathway framework, while Section 4.2 addresses the broader methodological implications of HYPER-NESS for hyperscanning research.

### 4.1 Empirical Support for the Dual-Pathway Model

#### 4.1.1 Neural–behavioral associations: functional differentiation of the two pathways

In the GF task (cooperative context) at ∼0.144 Hz, inter-brain contributions in LL and LH dyads showed comparable magnitudes but distinct behavioral associations. In LL dyads, inter-brain contribution was positively associated with attentional bias toward game rewards, whereas in LH dyads it was negatively associated with attentional bias toward social rewards. This dissociation indicates that equivalent levels of inter-brain coordination may relate to different reward domains depending on dyadic composition.

Within the proposed framework, the LL pattern is broadly consistent with the content resonance account, in which interpersonal coordination aligns with enhanced sensitivity to task-relevant rewards (Jackson et al., 2017; Smaldino, 2023). The LH pattern is consistent with the social contagion account, in which coordination is associated with reduced sensitivity to social reward cues — a feature commonly reported in individuals with higher levels of gaming addiction (Rosendo-Rios et al., 2022; Pastor-Satorras et al., 2015). Importantly, this distinction emerges at the level of functional association rather than the magnitude of inter-brain contribution itself.

At ∼0.02 Hz, neural–behavioral associations were observed only in the SF task (division-of-labor context). In LL dyads, intra-brain contribution was negatively associated with social reward bias, whereas in LH dyads inter-brain contribution was negatively associated with game reward bias. The absence of significant associations at this frequency in the GF task suggests that slow-timescale dynamics may become more behaviorally informative when real-time coordination demands are reduced, allowing reward-related neural fluctuations to more directly predict behavioral reward sensitivity.

#### 4.1.2 Multiscale causal structure: pathway differentiation from slow to fast dynamics

The neural–behavioral patterns described above are embedded within a multiscale temporal structure. At ∼0.144 Hz, both LL and LH dyads exhibited bidirectional directed temporal dependencies between inter-brain and intra-brain contributions across both tasks and both individuals within the dyad. This shared bidirectional structure suggests that reciprocal coupling between individual neural activity and inter-brain coordination at faster timescales may reflect a general property of interpersonal coordination rather than a pathway-specific feature (Sebanz et al., 2006; Hasson et al., 2012).

Differentiation between dyad types emerged at the slower ∼0.02 Hz band. In the GF task, LL dyads showed a significant unidirectional influence from intra-brain to inter-brain contribution (IntraL → Inter), whereas neither direction reached significance in LH dyads. This presence-versus-absence contrast suggests that, in LL dyads, internally generated reward-related dynamics showed predictive temporal precedence over interpersonal coordination, whereas this predictive relationship was absent in LH dyads.

In the SF task, both groups showed a significant inter-brain to intra-brain influence (Inter → IntraL). In LH dyads, an additional directed temporal dependency from the high-addiction individual’s intra-brain contribution to the inter-brain channel was observed (IntraH → Inter). This additional pathway, absent in LL dyads, is consistent with a more asymmetric coupling structure in LH dyads, in which neural dynamics originating from the high-addiction individual may propagate through the inter-brain channel to the low-addiction partner.

#### 4.1.3 Integrated interpretation: the complete mechanism of the dual pathways

Taken together, the three levels of evidence — neural–behavioral associations, Granger causality, and cross-frequency coupling — converge on a distinction in how slow reward dynamics and fast interpersonal coordination are coupled at the intra-brain/inter-brain interface across the two groups.

In LL dyads, this coupling appears reciprocal and symmetric. Individual reward-related dynamics contribute to driving inter-brain coordination (IntraL → Inter at ∼0.02 Hz), slow oscillations modulate fast coordination in the partner via cross-frequency coupling, and stronger inter-brain coordination is associated with heightened game reward sensitivity. Within the content resonance framework, these patterns are consistent with a self-reinforcing process in which addiction-related motivation may emerge from the dyadic interaction itself rather than from a single dominant source (Jackson et al., 2017; Smaldino, 2023; Bandura, 1977).

In LH dyads, the symmetric structure is replaced by a more asymmetric organization. The high-addiction individual’s internal dynamics contribute to the inter-brain channel (IntraH → Inter), which in turn is associated with changes in the low-addiction partner’s internal processing (Inter→ IntraL). The proactive intra-to-inter contribution of the low-addiction individual is absent, and no significant cross-frequency coupling was observed in either direction. At the behavioral level, stronger inter-brain coordination is associated with reduced social reward sensitivity in the low-addiction individual. This asymmetric pattern is broadly consistent with interpersonal neural entrainment frameworks, in which sequential coupling from one partner’s neural dynamics can shape the other’s processing in the absence of explicit behavioral mediation, with directionality tending to favor the more dominant partner (Wass et al., 2018, 2020). Within a putative social contagion framework, sustained exposure to a high-addiction partner’s reward-dominated neural dynamics may gradually increase the relative salience of game reward signals in the low-addiction individual’s processing, progressively compressing social reward sensitivity.

These findings speak to the theoretical gap identified by Van den Ende et al. (2022): integrating individual neuropsychological processes with social contagion perspectives requires analytical tools capable of simultaneously capturing interpersonal coordination and individual reward dynamics at the neural level. By decomposing interactive neural signals into inter-brain and intra-brain contributions, HYPER-NESS provides an analytical tool for such integration, revealing that the functional meaning and temporal organization of inter-brain coordination may differ systematically depending on the addiction composition of the dyad.

### 4.2 HYPER-NESS as a Methodological Advance in Hyperscanning Research

The present findings highlight a more general methodological limitation in hyperscanning research: conventional approaches do not provide the analytical solution required to disentangle distinct sources of inter-individual neural coordination. In this sense, the contribution of HYPER-NESS is not limited to the current application, but speaks to a broader issue in how inter-brain dynamics are traditionally quantified and interpreted.

First, as an extension of the FREQ-NESS framework (Rosso et al., 2025a), HYPER-NESS provides a principled, data-driven approach to estimating frequency-specific brain networks without imposing a priori assumptions on regions of interest. This is particularly relevant for fNIRS research, where the functional organization of brain activity is strongly shaped by slow, frequency-dependent hemodynamic fluctuations. Building on the work of Sasai et al. (2011), which established frequency as a meaningful domain for functional connectivity in fNIRS signals, the present framework enables the decomposition of whole-brain activity into frequency-specific components in an unbiased manner. Importantly, it also yields both spatial patterns and corresponding activation time series, allowing these components to be directly interrogated in subsequent analyses.

Second, by extending this framework to dyadic hyperscanning data, HYPER-NESS introduces a methodological distinction that conventional hyperscanning analyses cannot readily support. Rather than treating inter-brain synchrony as a unitary construct, the method explicitly separates inter-brain and intra-brain contributions within a unified analytical space. This distinction is critical, because it allows inter-individual coordination to be studied as a structured system composed of multiple interacting sources, rather than as a single aggregated index.

A recurring challenge in hyperscanning research — noted by Burgess (2013) and elaborated by Hamilton (2021), Holroyd (2022), and Rosso et al. (2025b) — is that standard inter-brain synchrony (IBS) measures conflate two conceptually distinct sources of inter-individual neural coordination. These are a genuine interactive coupling arising from mutual influence between participants, and parallel individual responses to shared stimuli that produce correlated signals without any direct brain-to-brain interaction. This confound is structural rather than incidental: metrics such as wavelet coherence, PLV, and WPLI quantify co-fluctuation between two signals without distinguishing whether that co-fluctuation reflects interpersonal coupling or common environmental entrainment. As the contrast between PLV and WPLI results in recent dual-brain EEG work illustrates (Yang et al., 2026), different synchrony metrics can yield divergent conclusions depending on their sensitivity to zero-lag versus lagged coupling, yet neither resolves the more fundamental question of whether the observed coordination is interactive in origin. Network-level approaches have partially addressed the related problem of regional specificity — for instance, Li et al. (2026) showed that combining time-domain IBS with spatial inter-subject similarity across distributed subnetworks predicted dyadic decision-making behavior more robustly than either measure alone — but decomposition in these frameworks is still performed per individual, leaving inter-brain and intra-brain contributions aggregated in the final synchrony index.

HYPER-NESS addresses this by performing GED on the concatenated dual-brain data matrix, yielding variance contributions that are inter-brain and intra-brain in origin, expressed in the same units, and directly comparable within the same analytical framework. The practical consequence for the present results is that LL and LH dyads were found to differ not in the magnitude of their inter-brain contributions but in the behavioral correlates of those contributions. At 0.144 Hz during GF task, associated with game reward attention bias in LL dyads (positively) and with social reward attention bias in LH dyads (negatively). This pattern is not detectable by any measure that treats inter-brain synchrony as a single quantity, because it requires correlating inter-brain and intra-brain contributions separately with independent behavioral measures. Standard IBS indices, which aggregate these two sources of variance, do not permit this differentiation.

The same point applies to the Granger causality analyses. Detecting the presence versus absence of IntraL→Inter directionality across groups required not only separating inter-brain from intra-brain contributions, but recovering their respective time series. Li et al. (2026) explicitly identified causal modeling as a direction for future work, noting that their framework did not support conclusions about directed temporal dynamics between inter-brain and intra-brain levels. The present pipeline addresses this by recovering activation time series for each source separately, allowing their temporal ordering to be examined. That the directionality of inter–intra coupling differs between groups independently of how much inter-brain contribution is present is a finding that presupposes this kind of decomposition.

More broadly, these results suggest that inter-brain synchrony magnitude and the functional meaning of that synchrony are dissociable quantities, and that studying their relationship requires methods that can separate inter-brain from intra-brain contributions rather than conflating them. This distinction is likely to be relevant beyond the present application: in any hyperscanning context where the theoretical question concerns whether neural co-fluctuation reflects genuine dyadic interaction or parallel responses to shared environmental input, treating inter-brain synchrony as a unitary construct may obscure the very effect under investigation.

## Supporting information

Table S1 (Component2)

## Acknowledgements

The Center for Music in the Brain (MIB) is funded by the Danish National Research Foundation (project number DNRF117), The Lundbeck Foundation (R469-2024-1573) and Købmand Herman Sallings Fond.

L.B. is supported by Sapere Aude: Independent Research Fund Denmark (DFF) Research Leader (grant ID: 10.46540/5253-00003B), Center for Music in the Brain, Linacre College of the University of Oxford.

M.R. is supported by Center for Music in the Brain and Nordic Mensa Fund.

