## Appendix A for "Gaming Addiction Transmission: Interpersonal Neural Pathways Revealed by HYPER-NESS"

**A.1 Time Series Recovery and Preprocessing**

Instantaneous contribution time series were constructed as described in Section 2.4.2.2: inter-brain contributions as the cross-product of the two participants' filtered signals, Inter(*t*) = *s*₁(*t*) × *s*₂(*t*), and intra-brain contributions as instantaneous power, Intra(*t*) = *s*(*t*)². Although the carrier frequencies are slow (C1: 0.020–0.026 Hz; C2: 0.144–0.148 Hz), power spectral analysis of the resulting envelope time series revealed faster fluctuations with a peak near 0.25 Hz, likely reflecting discrete game events (e.g., kill occurrences with ~4 s transient responses) modulating the slow-wave carrier amplitude. This faster envelope dynamics justified applying VAR-based Granger causality to the instantaneous power time series rather than to the raw carrier signals. Prior to model fitting, all time series were log-transformed to stabilize variance, linearly detrended to remove slow drift, and z-score standardized.

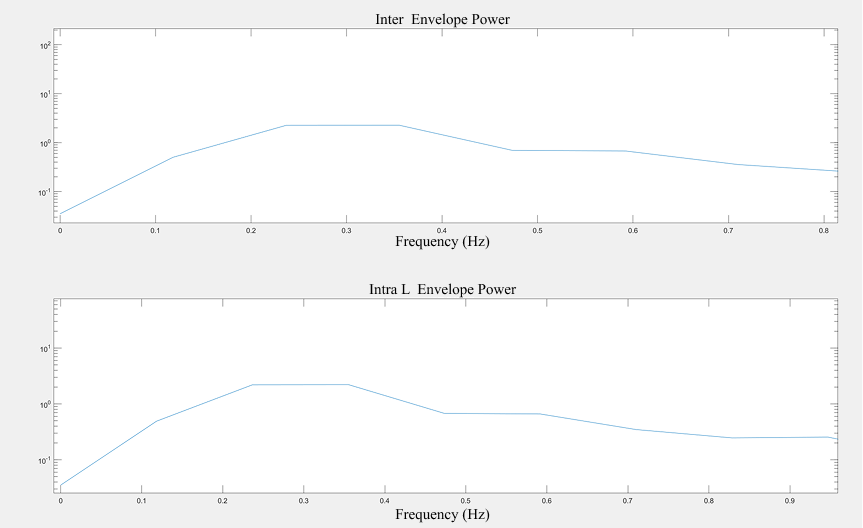

**A.2 Model Order Selection**

VAR model order was selected per dyad using the Akaike Information Criterion (AIC). Across dyads and conditions, AIC-selected orders were predominantly p = 1–2, indicating that immediate temporal dependencies dominated the envelope dynamics and that higher-order lags did not contribute meaningful predictive information beyond measurement noise. Based on these results, a fixed lag = 4 was adopted for the main analysis. The choice of lag = 4 corresponds to a temporal window of approximately 132 ms at the 30.3 Hz fNIRS sampling rate, which is sufficient to capture the rapid event-driven modulations identified in the envelope spectrum while avoiding overfitting given the relatively short task segments (~30 s, ~850 samples).

**A.3 Permutation Scheme**

Permutation testing was conducted with direction-specific asymmetric shuffling to preserve the autocorrelation structure of the predictor while disrupting only the directional predictive relationship of interest. Specifically, for the direction X → Y, the predicted variable Y was randomly permuted while the predictor X was kept intact; for the reverse direction Y → X, X was permuted while Y was kept intact. This asymmetric scheme ensures that the null distribution for each direction reflects the absence of that specific directed influence, rather than the absence of any temporal structure. Each direction was tested with 1,000 permutations, and p-values were computed as p = (# null ≥ observed + 1) / (n_perm + 1).

**A.4 Cross-Lag Consistency (Lag = 3–5)**

To verify that the main GC results were not sensitive to the specific lag choice, analyses were repeated at lag = 3 and lag = 5. Across both frequency clusters and both task conditions, the directional significance pattern was fully consistent across lags, with the following exception: in Cluster 2, LH Task 1, IntraH → Inter did not showed significance at lag = 3 for the 0.144 Hz frequency point (GC = 0.211, p = 0.211), but reaching significance at lag = 4 and lag = 5. This boundary attenuation is attributable to filter roll-off at the spectral edge of the cluster and does not affect the overall pattern. Quantitative GC values showed minor variation across lags consistent with the expected decrease in residual variance reduction beyond the AIC-selected order, but no reversal of directionality or change in statistical significance was otherwise observed (see figures and tables below for details).

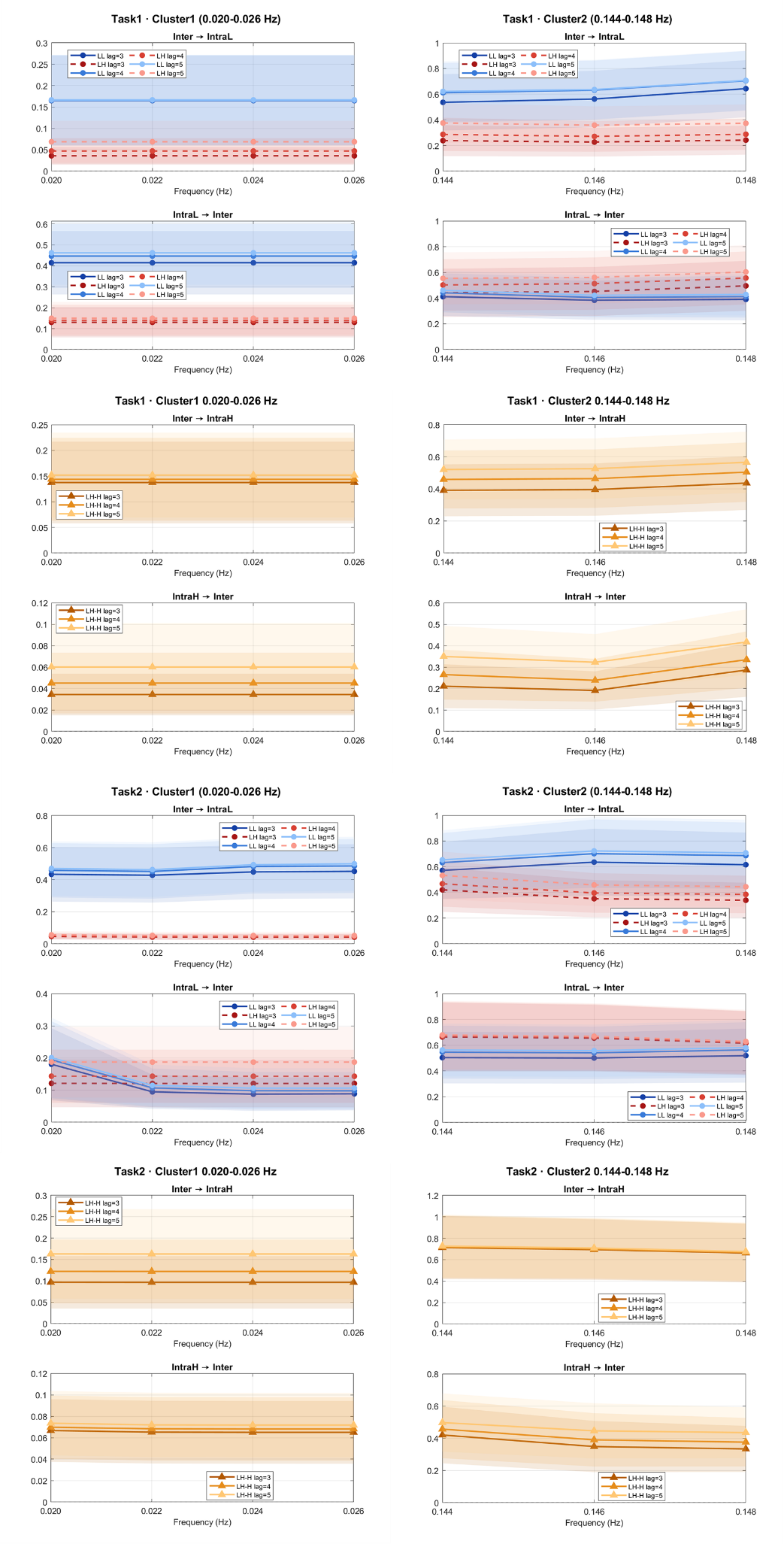

Cluster 1 (0.020–0.026 Hz) · Inter → IntraL

| **Freq** | LL Task1 | | | LL Task2 | | | LH Task1 | | | LH Task2 | | |
| --- | --- | --- | --- | --- | --- | --- | --- | --- | --- | --- | --- | --- |
|  | lag3 | lag4 | lag5 | lag3 | lag4 | lag5 | lag3 | lag4 | lag5 | lag3 | lag4 | lag5 |
| **0.02** | 0.137 | 0.136 | 0.134 | 0.021* | 0.016* | 0.014* | 0.104 | 0.135 | 0.179 | 0.024* | 0.024* | 0.026* |
| **0.022** | 0.137 | 0.136 | 0.134 | 0.023* | 0.017* | 0.015* | 0.104 | 0.135 | 0.179 | 0.032* | 0.032* | 0.035* |
| **0.024** | 0.137 | 0.136 | 0.134 | 0.017* | 0.011* | 0.010* | 0.104 | 0.135 | 0.179 | 0.035* | 0.034* | 0.037* |
| **0.026** | 0.137 | 0.136 | 0.134 | 0.017* | 0.011* | 0.009* | 0.104 | 0.135 | 0.179 | 0.035* | 0.034* | 0.037* |

Cluster 1 (0.020–0.026 Hz) · IntraL → Inter

| **Freq** | LL Task1 | | | LL Task2 | | | LH Task1 | | | LH Task2 | | |
| --- | --- | --- | --- | --- | --- | --- | --- | --- | --- | --- | --- | --- |
|  | l3 | l4 | l5 | l3 | l4 | l5 | l3 | l4 | l5 | l3 | l4 | l5 |
| **0.02** | 0.015* | 0.010* | 0.008** | 0.127 | 0.125 | 0.126 | 0.09 | 0.082 | 0.073 | 0.123 | 0.103 | 0.109 |
| **0.022** | 0.015* | 0.010* | 0.008** | 0.093 | 0.095 | 0.093 | 0.09 | 0.082 | 0.073 | 0.124 | 0.103 | 0.109 |
| **0.024** | 0.015* | 0.010* | 0.008** | 0.11 | 0.112 | 0.105 | 0.09 | 0.082 | 0.073 | 0.124 | 0.103 | 0.109 |
| **0.026** | 0.015* | 0.010* | 0.008** | 0.105 | 0.113 | 0.104 | 0.09 | 0.082 | 0.073 | 0.124 | 0.103 | 0.109 |

Cluster 2 (0.144–0.148 Hz) · Inter → IntraL

| **Freq** | LL Task1 | | | LL Task2 | | | LH Task1 | | | LH Task2 | | |
| --- | --- | --- | --- | --- | --- | --- | --- | --- | --- | --- | --- | --- |
|  | l3 | l4 | l5 | l3 | l4 | l5 | l3 | l4 | l5 | l3 | l4 | l5 |
| **0.144** | 0.026* | 0.018* | 0.018* | 0.020* | 0.014* | 0.012* | 0.065 | 0.042* | 0.027* | 0.026* | 0.018* | 0.011* |
| **0.146** | 0.021* | 0.015* | 0.014* | 0.026* | 0.018* | 0.015* | 0.061 | 0.038* | 0.026* | 0.028* | 0.021* | 0.013* |
| **0.148** | 0.011* | 0.008** | 0.008** | 0.025* | 0.017* | 0.014* | 0.049* | 0.031* | 0.022* | 0.024* | 0.018* | 0.011* |

Cluster 2 (0.144–0.148 Hz) · IntraL → Inter

| **Freq** | LL Task1 | | | LL Task2 | | | LH Task1 | | | LH Task2 | | |
| --- | --- | --- | --- | --- | --- | --- | --- | --- | --- | --- | --- | --- |
|  | l3 | l4 | l5 | l3 | l4 | l5 | l3 | l4 | l5 | l3 | l4 | l5 |
| **0.144** | 0.015* | 0.010* | 0.008** | 0.023* | 0.017* | 0.014* | 0.031* | 0.023* | 0.015* | 0.024* | 0.023* | 0.021* |
| **0.146** | 0.022* | 0.017* | 0.014* | 0.023* | 0.017* | 0.014* | 0.031* | 0.023* | 0.023* | 0.022* | 0.021* | 0.019* |
| **0.148** | 0.029* | 0.023* | 0.018* | 0.025* | 0.019* | 0.015* | 0.021* | 0.016* | 0.010* | 0.024* | 0.023* | 0.024* |

Cluster 1 (0.020–0.026 Hz) · IntraH → Inter

| **Freq** | LH_Task1 | | | LH_Task2 | | |
| --- | --- | --- | --- | --- | --- | --- |
|  | lag=3 | lag=4 | lag=5 | lag=3 | lag=4 | lag=5 |
| **0.02** | 0.034 | 0.045 | 0.06 | 0.067* | 0.070* | 0.074* |
| **0.022** | 0.034 | 0.045 | 0.06 | 0.065* | 0.068* | 0.072* |
| **0.024** | 0.034 | 0.045 | 0.06 | 0.065* | 0.068* | 0.072* |
| **0.026** | 0.034 | 0.045 | 0.06 | 0.065* | 0.068* | 0.072* |

Cluster 1 (0.020–0.026 Hz) · Inter → IntraH

| **Freq** | LH_Task1 | | | LH_Task2 | | |
| --- | --- | --- | --- | --- | --- | --- |
|  | lag=3 | lag=4 | lag=5 | lag=3 | lag=4 | lag=5 |
| **0.02** | 0.137 | 0.144 | 0.152 | 0.097 | 0.122 | 0.163 |
| **0.022** | 0.137 | 0.144 | 0.152 | 0.097 | 0.122 | 0.163 |
| **0.024** | 0.137 | 0.144 | 0.152 | 0.097 | 0.122 | 0.163 |
| **0.026** | 0.137 | 0.144 | 0.152 | 0.097 | 0.122 | 0.163 |

Cluster 2 (0.144–0.148 Hz) · Inter → IntraH

| **Freq** | LH_Task1 | | | LH_Task2 | | |
| --- | --- | --- | --- | --- | --- | --- |
|  | lag=3 | lag=4 | lag=5 | lag=3 | lag=4 | lag=5 |
| **0.144** | 0.391* | 0.459* | 0.521* | 0.713* | 0.719* | 0.727* |
| **0.146** | 0.396* | 0.464* | 0.526* | 0.694* | 0.701* | 0.709* |
| **0.148** | 0.436* | 0.504* | 0.566** | 0.662* | 0.667* | 0.676* |

Cluster 2 (0.144–0.148 Hz) · IntraH → Inter

| **Freq** | LH_Task1 | | | LH_Task2 | | |
| --- | --- | --- | --- | --- | --- | --- |
|  | lag=3 | lag=4 | lag=5 | lag=3 | lag=4 | lag=5 |
| **0.144** | 0.211 | 0.265* | 0.350* | 0.420* | 0.457* | 0.497* |
| **0.146** | 0.191* | 0.239* | 0.323* | 0.348* | 0.390* | 0.446* |
| **0.148** | 0.287* | 0.335* | 0.417* | 0.334* | 0.376* | 0.435* |

**A.5 Directionality Interpretation**

A critical interpretive principle governing the GC results is that the same directed relationship cannot simultaneously reflect active influence in one group and passive tracking in another. GC direction is a fixed property of the temporal precedence structure: if Inter(t) Granger-causes IntraL(t), this means that past values of the inter-brain signal carry predictive information about future intra-brain activity, regardless of group membership. The group difference lies not in the direction itself but in whether the reverse direction is also present. In LL dyads, bidirectional GC (Inter → IntraL and IntraL → Inter) indicates a closed-loop mutual modulation consistent with active neural coordination between partners. In LH dyads, the presence of IntraL → Inter without the reverse direction indicates that the individual's intra-brain dynamics systematically precede and predict the inter-brain coupling pattern, without reciprocal modulation from the inter-brain level — a pattern consistent with passive contagion tracking rather than active bidirectional coordination.
