## Appendix B for "Gaming Addiction Transmission: Interpersonal Neural Pathways Revealed by HYPER-NESS"

**B.1 Cross-Frequency Coupling Analysis**

Phase-amplitude coupling (PAC) between the two identified frequency clusters (C1: 0.020–0.026 Hz; C2: 0.144–0.148 Hz) was quantified using the modulation index (MI) proposed by Tort et al. (2010), defined as the Kullback-Leibler divergence between the observed amplitude distribution over phase bins and a uniform distribution:

$$\text{MI}=\frac{\sum_{j} P(j)\log[P(j)\cdot n_{\text{bins}}]}{\log(n_{\text{bins}})}$$

where *P*(*j*) is the mean amplitude in the *j*-th phase bin normalized to sum to one, and *n*_bins = 18. Time series for CFC were constructed identically to the Granger causality analysis (see Appendix A.1). Phase was extracted from the filtered signal via the Hilbert transform; amplitude was operationalized as instantaneous power (squared filtered signal).

A representative frequency pair was selected from each cluster (C1 = 0.020 Hz, C2 = 0.144 Hz) for the main analysis. Group-level statistical inference was conducted via permutation testing (*n* = 1,000): on each iteration, the phase time series of each dyad was independently circularly shifted by a random lag, PAC was recomputed, and the group mean MI was recorded to form a null distribution. Observed group mean MI was compared against this null distribution as *p* = (# null ≥ observed + 1) / (*n*_perm + 1). For cross-subject CFC, phase and amplitude were drawn from individual-level signals; in LL dyads the symmetric mean of both cross-subject directions served as a single undirected index, while in LH dyads directions were remapped according to addiction group membership (H→L and L→H separately). A mixed ANOVA (Group × Task) was additionally conducted on cross-subject MI values.

**B.2 Robustness Analysis Across Frequency-Pair Combinations**

Robustness analysis across all 12 frequency-pair combinations (C1: 0.020–0.026 Hz × C2: 0.144–0.148 Hz) confirmed the specificity and directionality of the main finding.

For the C1ph→C2amp direction, the LL_GF (task 1) condition was the only condition to show significant coupling: 9 of 12 combinations reached significance (permutation test, p < 0.05), with highly consistent MI values across the frequency grid (range: 0.126–0.136; CV < 3%). The three non-significant combinations all involved C1 = 0.026 Hz, the spectral boundary of the slow-wave cluster where filter attenuation is expected given the Gaussian filter FWHM. In all other conditions (LL_SF (task 2), LH_GF (task 1), LH_SF (task 2)), 0 of 12 combinations reached significance, with MI values consistently lower than LL_GF (task 1) (approximately 0.105–0.109 vs. 0.129–0.136). In the LH group, MI values were identical across all four C1 frequency points, confirming the expected spectral redundancy within the slow-wave cluster due to filter overlap.

Together, these results demonstrate that the cross-subject CFC effect is specific to LL dyads during cooperative gaming (Task 1) and robust to the selection of representative frequency points within both clusters.

**LL Dyads**

|  | **LL — GF Task (T1)** | | | **LL — SF Task (T2)** | | |
| --- | --- | --- | --- | --- | --- | --- |
| **C1 (Hz)** | **C2=0.144** | **C2=0.146** | **C2=0.148** | **C2=0.144** | **C2=0.146** | **C2=0.148** |
| 0.020 | **0.1357*** | **0.1359*** | **0.1333*** | 0.1293 | 0.1293 | 0.1293 |
| 0.022 | **0.1288*** | **0.1289*** | **0.1264*** | 0.1292 | 0.1293 | 0.1293 |
| 0.024 | **0.1287*** | **0.1289*** | **0.1263*** | 0.1294 | 0.1295 | 0.1295 |
| 0.026 | 0.1292 | 0.1294 | 0.1268 | 0.1300 | 0.1301 | 0.1301 |

**LH Dyads — GF Task (T1)**

|  | **LH, GF Task (T1): H→L** | | | **LH, GF Task (T1): L→H** | | |
| --- | --- | --- | --- | --- | --- | --- |
| **C1 (Hz)** | **C2=0.144** | **C2=0.146** | **C2=0.148** | **C2=0.144** | **C2=0.146** | **C2=0.148** |
| 0.020 | 0.1088 | 0.1049 | 0.1044 | 0.1094 | 0.1039 | 0.1032 |
| 0.022 | 0.1088 | 0.1049 | 0.1044 | 0.1094 | 0.1039 | 0.1032 |
| 0.024 | 0.1088 | 0.1049 | 0.1044 | 0.1094 | 0.1039 | 0.1032 |
| 0.026 | 0.1088 | 0.1049 | 0.1044 | 0.1094 | 0.1039 | 0.1032 |

**LH Dyads — SF Task (T2)**

|  | **LH, SF Task (T2): H→L** | | | **LH, SF Task (T2): L→H** | | |
| --- | --- | --- | --- | --- | --- | --- |
| **C1 (Hz)** | **C2=0.144** | **C2=0.146** | **C2=0.148** | **C2=0.144** | **C2=0.146** | **C2=0.148** |
| 0.020 | 0.1068 | 0.1067 | 0.1057 | 0.1078 | 0.1071 | 0.1079 |
| 0.022 | 0.1069 | 0.1068 | 0.1059 | 0.1076 | 0.1070 | 0.1076 |
| 0.024 | 0.1069 | 0.1068 | 0.1059 | 0.1076 | 0.1070 | 0.1076 |
| 0.026 | 0.1069 | 0.1068 | 0.1059 | 0.1076 | 0.1070 | 0.1076 |

Note: Modulation index (MI) values for all 12 frequency-pair combinations (C1: 0.020–0.026 Hz × C2: 0.144–0.148 Hz) across all conditions. Bold values with asterisk (*) indicate p < 0.05 (permutation test). Green shading marks significant cells. Identical MI values across C1 frequency points in LH dyads reflect expected spectral redundancy due to Gaussian filter FWHM exceeding the frequency spacing within C1. The three non-significant cells in LL GF Task (T1) all involve C1 = 0.026 Hz, the spectral boundary of the slow-wave cluster where filter attenuation is expected.

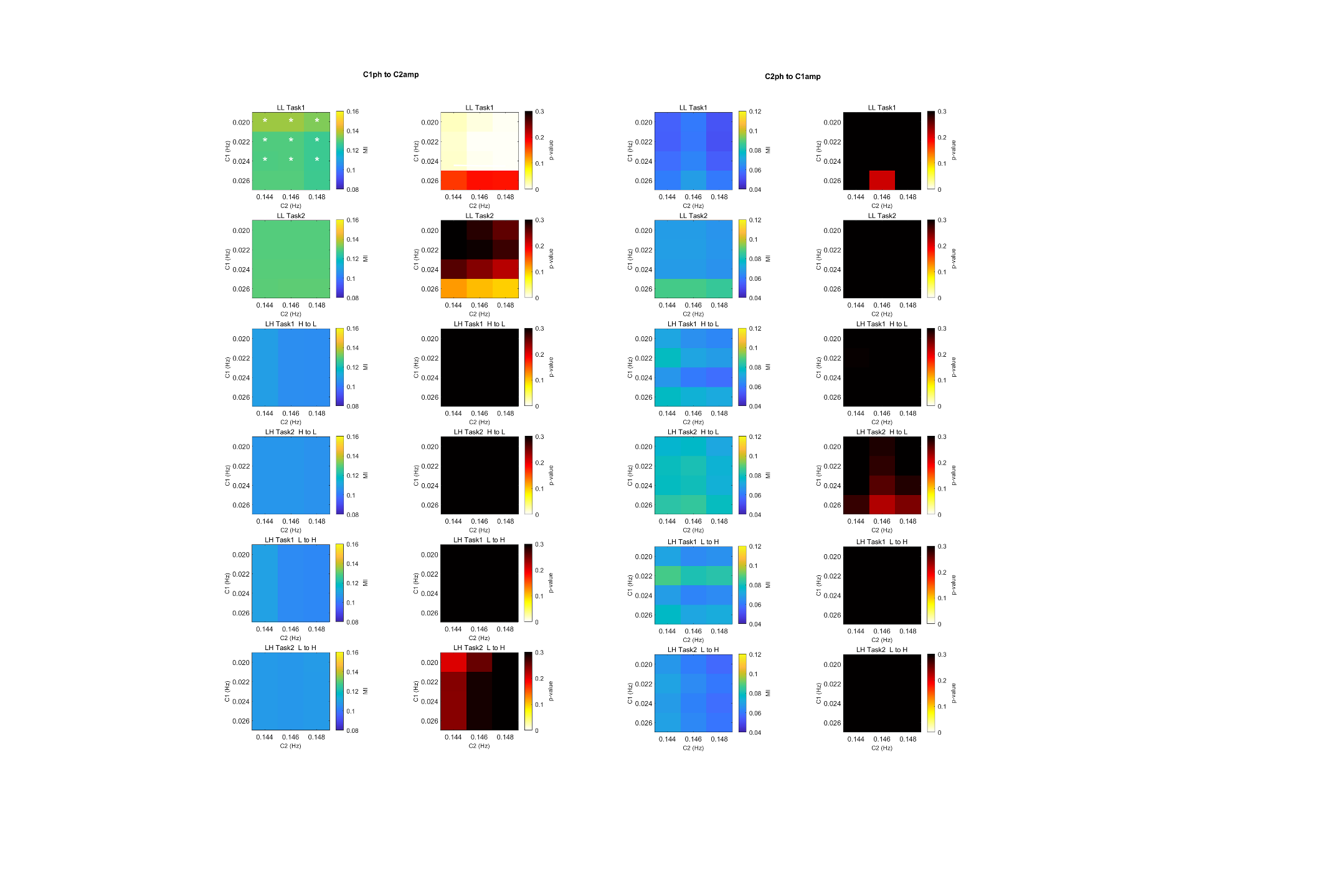
